# Multiple memory systems for efficient temporal order memory

**DOI:** 10.1101/2022.06.10.495702

**Authors:** Anna Jafarpour, Jack J. Lin, Robert T. Knight, Elizabeth A. Buffalo

## Abstract

We report the distinct contribution of multiple memory systems to retrieval of the temporal order of events. The neural dynamics related to retrieval of movie scenes revealed that recalling the temporal order of close events elevates hippocampal theta power, like that observed for recalling close spatial relationships. In contrast, recalling far events elevates beta power in the orbitofrontal cortex, reflecting recall based on the overall movie structure.

## Main text

Remembering the order of event occurrence is fundamental to episodic memory. Converging evidence suggests that at least two copies of events are encoded in parallel in distinct brain regions during memory formation^1–3^. One system retains events with high fidelity and is hippocampal-dependent. The other system encodes schematic information and engages the prefrontal cortex. At recall, memories are reconstructed when we combine our understanding of the unfolding of events in time with episodic information^4^. Episodic memory, especially recalling the temporal order of events, requires the hippocampal network, evident from the behavior of patients with hippocampal lesions who have impairments in recalling past events in the same order as they were encountered ^5,6^. Functional MRI and lesion studies suggest that in addition to the hippocampus (HPC), the orbitofrontal cortex (OFC) is involved in successful temporal order recall ^7,8^. However, the neural mechanisms by which the HPC and the OFC support temporal order memory are unclear.

Temporal memory, shaped by the relatedness of the sequence of events, can be studied as the absolute time relative to salient event boundaries ^9^ or as the recency of events relative to each other ^10,11^. At encoding, both the OFC and HPC track the temporal relatedness of events ^12^, and the left HPC is known to represent the temporal approximation of events ^13^. Temporal order recall is easier for events with distinct contexts that are most likely to occur far apart in time ^10^. This suggests that the temporal context of far events enables temporal order judgment without the need to recall the episodic details of an event’s recency. Recalling memories with high precision engages the HPC ^14^, however, tracking the temporal order of long sequences of events with high fidelity would require substantial HPC resources. Accordingly, we hypothesized that memory systems utilize OFC for recalling the temporal order of far events, while HPC retrieves the episodic details necessary for recalling the temporal order of close events.

We investigated the neural dynamics within the HPC and the medial OFC in the left hemisphere that support recognition of the temporal order of events from long-term memory. Epileptic patients undergoing seizure monitoring (n=8, 3 female, Table 1) watched a novel animated movie and, after brief delay including a 2-min rest period, determined the order of movie frames that were presented in pairs (Figure 1a and 1b). Subjects performed equally well in recalling close and far events (paired t-test t(7) = 1.7, p = 0.86). However, response times were longer for close events compared to far events (paired t-test t(7) = 2.64, p = 0.03; Figure 1c), indicating that temporal order judgement was more demanding for the close events.

**Figure 1.**
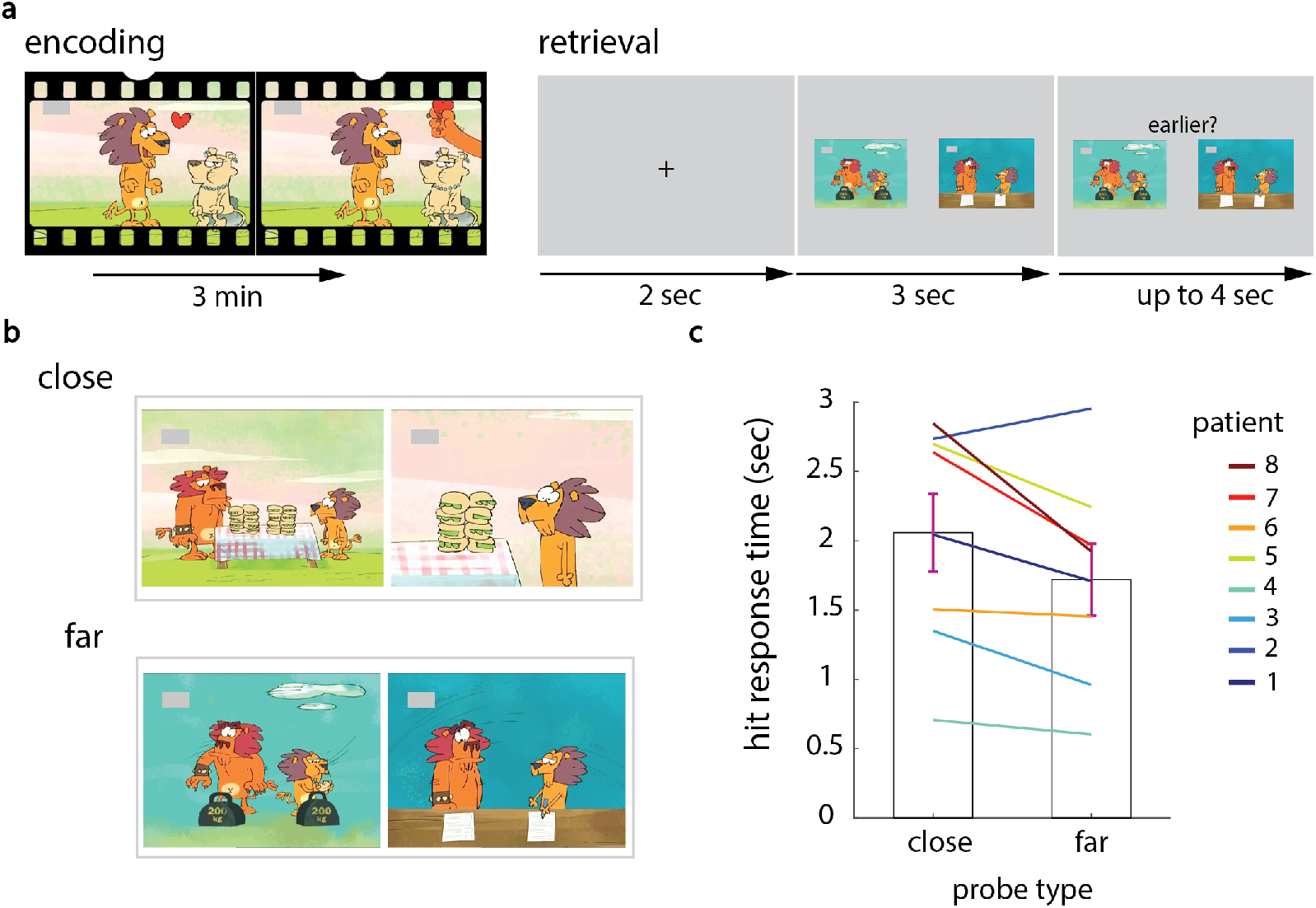
Experimental design and behavior. **(a)** At encoding, participants watched a novel animation (3 minutes long) and were informed that memory would be assessed. At retrieval, participants performed a temporal order recognition task. Each temporal order retrieval trial started with a fixation point to draw attention to the middle of the screen. Then, two frames of the movie were shown side by side. After 3 seconds, an instruction appeared asking about the order of the probes. Pseudo-randomly, the instruction was split to ask for selection of the earlier or later frame. This design allowed us to study the neural mechanism of memory recall that is separate from preparing for the response. **(b)** An example of the *close* (top) and *far* (bottom) prompts. The close probes were neighboring movie frames that were visually distinct, while the far probes were between 12 to 60 seconds apart and were more contextually distinct than the close events (see the supplemental methods). **(c)** It took longer to correctly recall the temporal order of close events than far events. Patients’ responses are color-coded and numbered according to (table S1). The error bars show S.E.M.

We compared the power spectral density (PSD) for correct temporal judgement of close and far events during the 2 seconds after the probe onset (i.e., prior to the cue for response selection). At a group-level analysis, we compared the coefficients of the contrast between close and far correct temporal judgments against zero (no difference) in a mixed effect model. The results showed that theta power was higher for recalling the order of close events than for far events in the HPC (n = 7; 4-8 Hz, peaking at 6.5 Hz, p < 0.001). By contrast, beta power in the left OFC was higher for recalling the order of far compared to close events (n = 6; 20-25 Hz, peaking at 22.5 Hz, p = 0.012; Figure 2). The results suggest enhanced theta/alpha-band power in the HPC is associated with successful retrieval of episodic information in temporal order recall, whereas enhanced beta-band power in the OFC supports the temporal order recall of far events.

**Figure 2.**
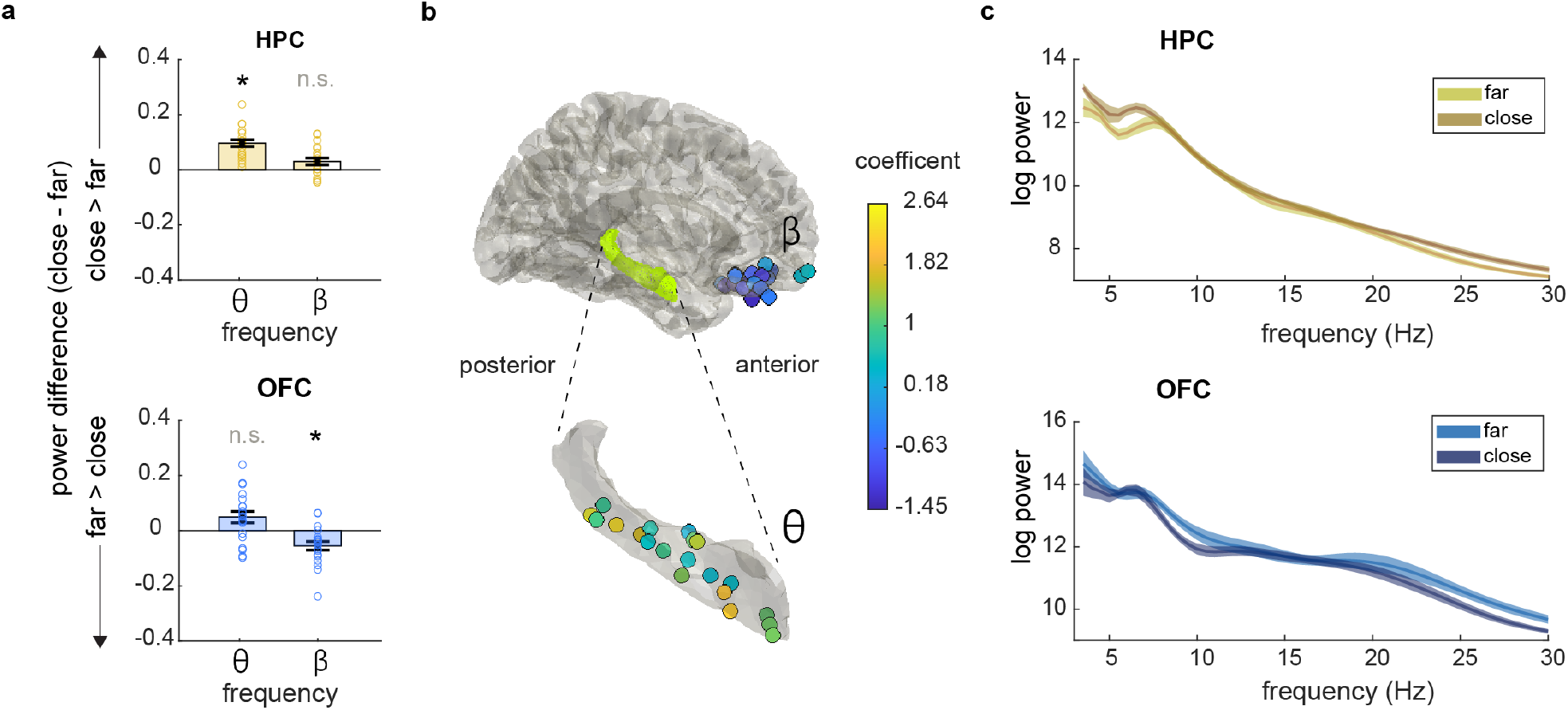
Hippocampal and OFC power differences between close and far events. **(a)** HPC theta-band (4-8 Hz) power was higher for close compared to far temporal memory recall (top). This effect was reversed for beta-band power in the OFC (20-25 Hz; bottom). * p < 0.005 and n.s. for p > 0.05. Error bars indicate S.E.M. **(b)** The coefficient for the comparison between recalling close and far events shown in the HPC (in theta-band) and in the OFC (in beta-band). A positive coefficient supports higher power in recalling close compared to far events; whereas a negative coefficient supports higher power in recalling far compared to close events. **(c)** Example of the PSD in one contact in the OFC (top) and one contact in the HPC (bottom). The shaded area shows S.E.M.

The hippocampus (HPC) is connected to the medial OFC both directly and indirectly through the entorhinal cortex and para-hippocampus gyrus ^15^. The entorhinal cortex - specifically the lateral entorhinal cortex along with the neighboring perirhinal cortex – has been implicated in the precision of recalling the time of an event relative to the task onset (a salient event boundary) ^9^ and these cortical regions have direct connections to both HPC and the OFC ^15^. Far events are more likely to have an intervening salient event boundary during encoding, raising the possibility that the order of far events can be inferred relative to the event boundaries and the gist of the movie structure without the need for hippocampal input. However, HPC is essential for temporal order memory of close events that are not anchored to the boundaries of a sequence ^16^. In the spatial domain, the absence of fine-resolution para-hippocampal representation disturbs the resolution of hippocampal spatial representation for places that are away from the salient boundaries ^17^. Similarly, we propose that, in parallel to the spatial neural mechanism, input from the entorhinal cortex to HPC supports recalling the temporal order of close events in fine detail. The OFC represents the gist structure of the flow of events that strengthens with consolidation ^18^, such as learning a reward value that follows an event ^19^. In contrast, HPC corresponds to the detailed order of events that closely follow each other ^20,21^. We predict that at retrieval, HPC provides the temporal order of close events following the top-down signal from the OFC, congruent with reports from rodents ^22^. Converging evidence suggests that the existence of multiple traces of memory benefits long-term memory consolidation ^1–3^. Here, we show that, in short time after encoding, the multiple memory traces in HPC and OFC support efficient temporal order memory, enabling recall of far events with the support of the OFC and close events with hippocampal activity.

## Methods and methods

The study protocol was approved by the Office for the Protection of Human Subjects of the University of California, Berkeley, the University of California, Irvine, and Stanford University. All patients provided written informed consent before participating and all methods were performed in accordance with the relevant guidelines and regulations.

### Participants

Eight participants (3 female, 5 right-handed, mean age = 38.8, sd = 14.3, range = 20-58) were implanted stereotactically with depth electrodes to localize the seizure onset zone for subsequent surgical resection (Table 1). All participants had normal or corrected normal vision. No seizures occurred during task administration. The signal for the epochs with interictal epileptiform activity was discarded. Only 3 of the participants had coverage for simultaneous recording from the general OFC and the HPC areas, which is too limited for conclusive across regions connectivity analysis.

| patient | Age (year) | gender | dominant hand | Number of contacts |  | Number of responded trials |  |  |
| --- | --- | --- | --- | --- | --- | --- | --- | --- |
|  |  |  |  | HPC | mOFC | Far | close | Hit rate (%) |
| 1 | 50 | M | R | 7 |  | 17 | 16 | 69 |
| 2 | 20 | M | R |  | 8 | 16 | 15 | 85 |
| 3 | 40 | F | L | 2 | 2 | 18 | 17 | 80 |
| 4 | 58 | F | R | 3 | 2 | 17 | 13 | 60 |
| 5 | 31 | M | L |  | 4 | 7 | 9 | 67 |
| 6 | 34 | M | R |  | 1 | 6 | 9 | 70 |
| 7 | 55 | F | L | 2 |  | 10 | 13 | 70 |
| 8 | 23 | M | R | 6 | 3 | 15 | 11 | 96 |

### Experimental design

Participants were instructed to carefully watch a short mute animation (around 3 minutes long) knowing that we would ask questions about the movie (Jafarpour et al, 2019 JoCN; link to the movie: https://www.youtube.com/watch?v=Q_guH9vA0sk). Then, after a brief delay of 3-5 minutes with distractions, the participants were instructed to perform a temporal order recognition test. This test required the participants to determine the order of two movie frames that were shown side by side. Each test trial started with a fixation point for two seconds, then two scenes of the movie were shown side by side for 3 sec. Right after, a probe appeared on top of the two frames, asking which frame occurred *earlier* or *later* (Figure 1). The probe was selected pseudo-randomly so that in half of the trials the earlier scene asked for the earlier scene. There was an equal number of trials with far and close probes. Participants used left or right arrow keys and had 4 sec available to respond. The experiment was designed to separate the process of retrieving the temporal order of events from picking an answer. The tested movie scenes were between 1.6 to 60 seconds apart allowing us to study the differences between temporal order recognition of events that were close in time and those that were far apart. The accuracy of temporal order recognition for three people who did not watch the movie and guessed the order showed that an accurate performance required watching the movie. The close probes were less than 11 seconds apart. An independent group of (n= 80) people determined the event boundaries in the movie, meaning when a new event started. The number of determined events in between the movie frames reflects the distinctness of the movie frames ^12^. In this task, the number of determined event boundaries between the close probes was significantly less than between the far probes (t(33) = 7.1, p < 0.001).

### EEG data collection and preprocessing

Intracranial EEG data were acquired using the Nihon Kohden recording system, analog-filtered above 0.01 Hz, and digitally sampled at 5 kHz or 10 kHz. A photodiode was recorded to track the flow of the experiment as it appeared to the participant. All EEG analyses were run in Matlab 2014a, EEGLAB, and Fieldtrip offline. Fieldtrip was used for electrode localization ^23^. The electrodes’ anatomical locations were determined by a neurologist’s team in the native space. The EEG recorded during the retrieval was de-trended (i.e., the mean value of the entire signal at each contact was subtracted for the value at each time point). Then the EEG was high-pass filtered at 0.5 Hz using a zero-phase delay finite impulse response filter with Hamming window, and then down-sampled to 1 Khz using Matlab’s resample() function. Independent component analysis of the signal of all recording channels with neuronal data was used for removing the common noise ^24^ – akin to a weighted common average re-referencing. The EEG was cropped around the onset of the probes when two movie frames were shown side by side with no instruction. Trials with the interictal epileptiform that a neurologist identified were discarded (Table for the number of trials per participant).

### Power Spectral Density

We contrasted the power spectrum density (PSD) of signals that were elicited by the far versus close probe onset. The power spectrum density was summarized after transformation of the signal round the probe onset into time and frequency 3 to 30 Hz with steps of 0.5 Hz using Morlet wavelet with 7 cycles. The power at every frequency was averaged for the first two seconds after the probe onset and contrasted between far and close trials using a linear regression. The coefficient of the contrast (Beta values) of each electrode in one region were pooled and averaged to have one value per region and patient for every frequency. The coefficients of the contrasts (beta-coefficients) of each electrode in one region were pooled. The aggregated coefficients were then compared against zero using a mixed-effect model with the patient number as a random factor. The p-values were adjusted to account for the False Discovery Rate (FDR) related to multiple comparisons of far and close trials at every frequency ^25^. We accepted p<0.025 because we repeated the comparison in two regions. We only reported the FDR-corrected p-values. Although the analysis is done in high frequency resolution, for visualization purposes, Figure 2 shows a summary of the relative power differences in a frequency band, formulated as (*close*−*far*)/(*close*+*far*).

## Notes

### Competing Interest Statement

The authors have declared no competing interest.

